# Structural insight into antibody evasion of SARS-CoV-2 omicron variant

**DOI:** 10.1101/2022.01.25.477671

**Authors:** Jyoti Verma, Naidu Subbarao

## Abstract

The severe acute respiratory syndrome coronavirus 2 (SARS-CoV-2) continues to mutate and evolve with the emergence of omicron (B.1.1.529) as the new variant of concern. The rapid spread of this variant regionally and globally could be an allusion to increased infectivity, transmissibility, and antibody resistance. The omicron variant has a large set of mutations in its spike protein, specifically in the receptor binding domain (RBD), reflecting their significance in ACE2 interaction and antibody recognition. We have carried out the present study to understand how these mutations structurally impact the binding of the antibodies to their target epitope. We have computationally evaluated the binding of different classes of RBD targeted antibodies, namely, CB6 (etesevimab), REGN10933 (casirivimab), S309 (sotrovimab), and S2X259 to the omicron mutation-induced RBD. Molecular dynamics simulations and binding free energy calculations unveil the binding affinity and stability of the antibody-RBD complexes. All the four antibodies show reduced binding affinity towards the omicron RBD. The therapeutic antibody CB6 aka etesevimab was substantially affected due to numerous omicron mutations occurring in its target epitope. This study provides a structural insight into the reduced efficacy of RBD targeting antibodies against the SARS-CoV-2 omicron variant.

## 1. Introduction

The continuous evolution of severe acute respiratory syndrome coronavirus 2 (SARS-CoV-2) has resulted in the resurgence of novel variants throughout the COVID-19 pandemic. The world health organization (WHO), on November 26, 2021, announced the addition of a new variant named Omicron (B.1.1.529) into the list of Variants of Concern (VOCs) [1]. The B.1.1.529 sequences were predominantly detected in South Africa, Botswana, and Hong Kong; however, it was first reported to the WHO from South Africa [1, 2]. As of January 25, 2022, the GISAID database has a collection of 588,320 omicron genome sequences shared from 125 countries, reflecting the rapid spread of the variant across the world [3]. The omicron variant has about 60 mutations compared to the reference SARS-CoV-2 genome, and many of these mutations have not been previously observed in other variants [4]. These mutations are present in the genomic regions encoding orf1a, orf1b, orf9b, spike protein, envelope protein, membrane protein, and nucleocapsid protein. However, a large number of mutations observed in the spike gene are of primary concern. There are 30 amino acid substitutions, one small insertion, and three deletions in the spike protein, out of which 15 are located in the receptor binding domain (RBD) (Figure 1). Thus, these variations may play an essential role in ACE2 binding and antibody recognition. A cluster of mutations (H655Y, N679K, and P681H) present at the furin cleavage site may be associated with enhanced transmissibility [5]. The substitutions Q498R and N501Y are reported to increase the binding affinity with ACE2 [6, 7]. There are also various mutations that are associated with the resistance to neutralizing and therapeutic monoclonal antibodies [8, 9]. Therefore, these mutations could effectively reduce the efficacy of the current vaccines and therapeutic antibodies against COVID-19. Scientists around the world have confirmed this threat by their findings. A preliminary report from Wilhelm et al. shows complete loss of neutralization of omicron strain in sera isolated from vaccinated individuals [10]. These observations were supported by another study where they report that B.1.1.529 is resistant to neutralization by serum from convalescent patients and individuals vaccinated and boosted with mRNA-based vaccines [11]. Studies also report the omicron variant’s immune escape and antibody resistance [10–13].

**Figure 1:**
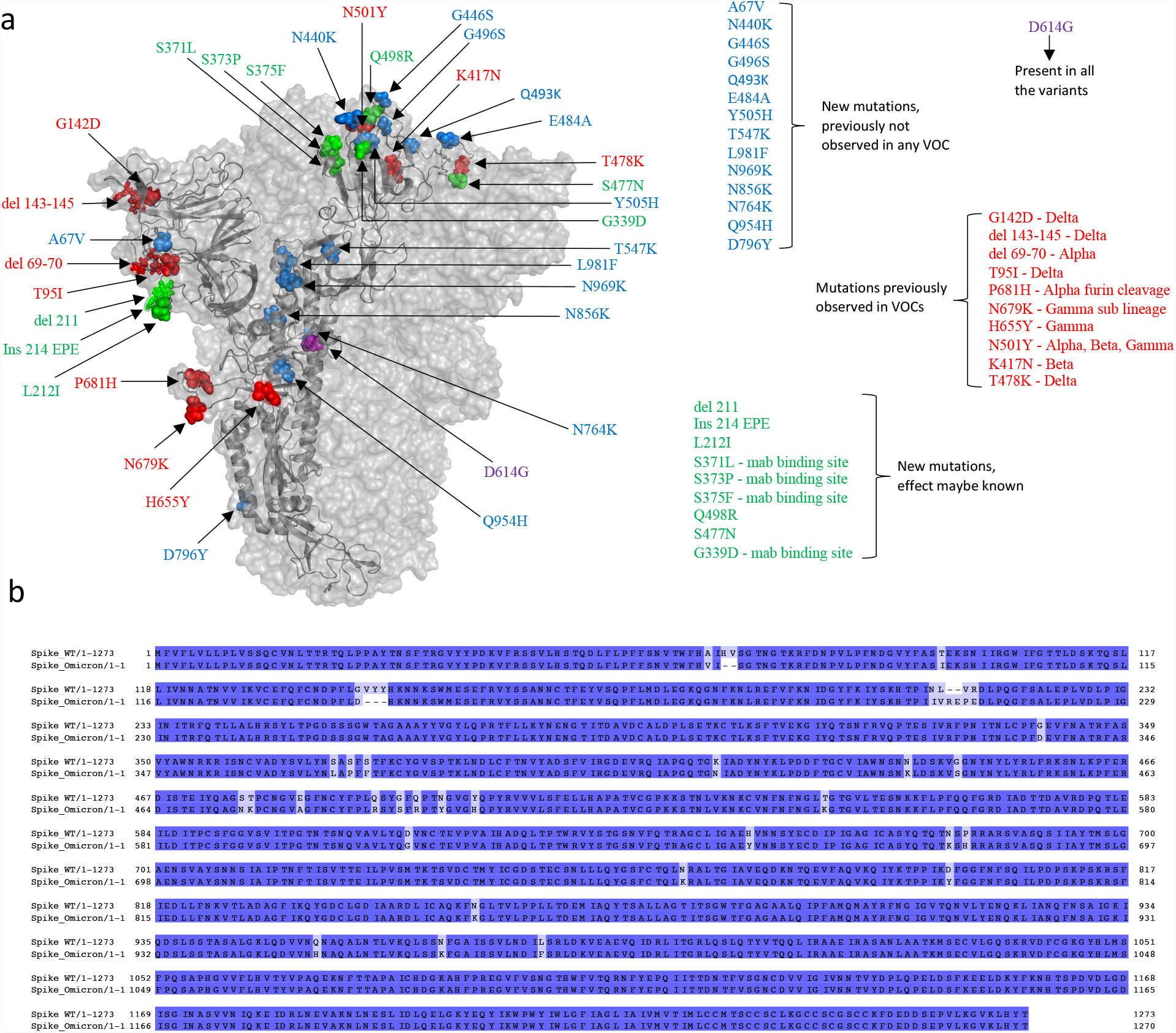
SARS-CoV-2 omicron mutations in the spike protein. (a) Spike protein mutation sites in omicron. The mutation sites are shown in spheres. (b) Pairwise sequence alignment of the wild type spike protein and omicron variant spike protein.

Thus, our study aims to understand the difference in binding of RBD targeted antibodies to the wildtype and omicron (B.1.1.529) RBD structure. In this study, we have carried out the structural evaluation of different class of antibodies in complex with the WT and omicron RBD. The RBD targeted neutralizing antibodies (NAbs) have been categorized into different classes based on their structural features and mode of action [14]. Class 1 NAbs (for ex., CB6 (etesevimab), Brii-196 (amubarvimab), 1-20, C102) primarily targets the receptor binding motif (RBM) and binds to the up conformation of the RBD to block ACE2 interaction [15–17]. Class 2 NAbs (REGN10933 (casirivimab), LY-CoV555 (bamlanivimab), COV2-2196) binds to the ACE2 site in both up and down RBD conformations [18–20]. Class 3 antibodies like S309 (sotrovimab) and REGN10987 (imdevimab) target the conserved core domain without altering ACE2 interactions [18, 21]. Antibodies from class 4 (DH1047, S2X259, ADG-2) targets epitopes spanning both the RBM and the core domain [22–24].

Here, we selected four different therapeutic antibodies, one from each class, and employed a computational approach to understand the energetics of their binding with the WT and omicron RBD using the Molecular dynamic simulations and binding free energy calculations.

## 2. Methodology

### 2.1 Data retrieval and structure preparation

We collected the sequence information for surface glycoprotein of SARS-CoV-2 wild type (WT) and the B.1.1.529 variant (Omicron) from the NCBI database with the GenBank IDs YP_009724390.1 and UFO69279.1, respectively [25, 26]. A pairwise alignment was performed for both the spike protein sequences using the EMBOSS Needle program [27]. Further, the structures of various antibodies bound to the spike protein were retrieved from the RCSB Protein Data Bank (http://www.rcsb.org/pdb/), and their epitope information was collected from the IEDB database (http://www.iedb.org/) [28, 29]. For our study, we selected CB6 (PDB ID: 7C01), REGN10933 (PDB ID:6XDG), S309 (PDB ID: 7R6X) and S2X259 (PDB ID: 7M7W) antibody complexes. The B.1.1.529 variant mutations were introduced in the selected RBD-antibody crystal structures with the maestro interface of the Schrodinger suite (v2019.1) [30]. These raw structures were further processed in the protein preparation wizard for the addition of missing amino acid side chains, addition of missing hydrogen atoms, and optimization of the hydrogen bond network. The structures were finally energy minimized with the OPLS3e force field and saved for further calculations.

### 2.2 Molecular Dynamics simulation

We carried out the MD simulation of the RBD-antibody structures and their respective mutant (including B.1.1.529 mutations) complexes. Hence, a total of 8 individual simulation systems were prepared with identical parameters. The simulation was performed with the GROMACS package v2019.4 [31]. The systems were parameterized with the AMBER99SB force field in a cubic box filled with TIP3P water molecules. Neutralization was achieved by adding the required number of counter ions (Na+/Cl−), and energy minimization was carried out using the steepest descent minimization algorithm. Before the final MD run, the systems were equilibrated in two consecutive steps, NVT (particles, volume, and temperature kept constant) and NPT (particles, pressure, and temperature kept constant) for 100 picoseconds (ps) each. Finally, each system was simulated for 100 nanoseconds (ns), and energies were saved every 10 ps.

### 2.3 Interaction analysis and binding free energy calculation

The MD simulation trajectory of each RBD-antibody complex was analyzed, and coordinates of the lowest energy conformation of the complex were extracted for further interaction analysis and energy calculations. We used the web application COCOMAPS (bioCOmplexes COntact MAPS) for evaluating the interface and interaction statistics of the predicted conformations of the RBD-antibody complexes [32, 33]. Further, we calculated the binding free energies of all the complexes with the Molecular Mechanics/Generalized Born Surface Area (MMGBSA) approach using the HawkDock server [34, 35].

## 3. Results and discussion

The B.1.1.529 spike protein mutations are spread throughout the RBD and the NTD, which are the main antigenic target regions of therapeutic or vaccine-generated antibodies. The majority of experimentally validated epitopes occur in the RBD of spike glycoprotein. Some of the antibodies with their structure and epitope information are mentioned in Table 1. We collected the SARS-CoV-2 specific spike protein antigen epitope assay data from the IEDB database. The number of positive epitope assays are indicated for each residue position in figure 2a, and the immune response frequency for each of the position in the spike antigen is shown in figure 2b. The B.1.1.529 mutation mapping revealed a large cluster of mutations occurring at the residue positions with higher immune response frequency. Hence, the question arises of how these mutations structurally impact the binding of the RBD targeted antibodies.

**Table 1:**
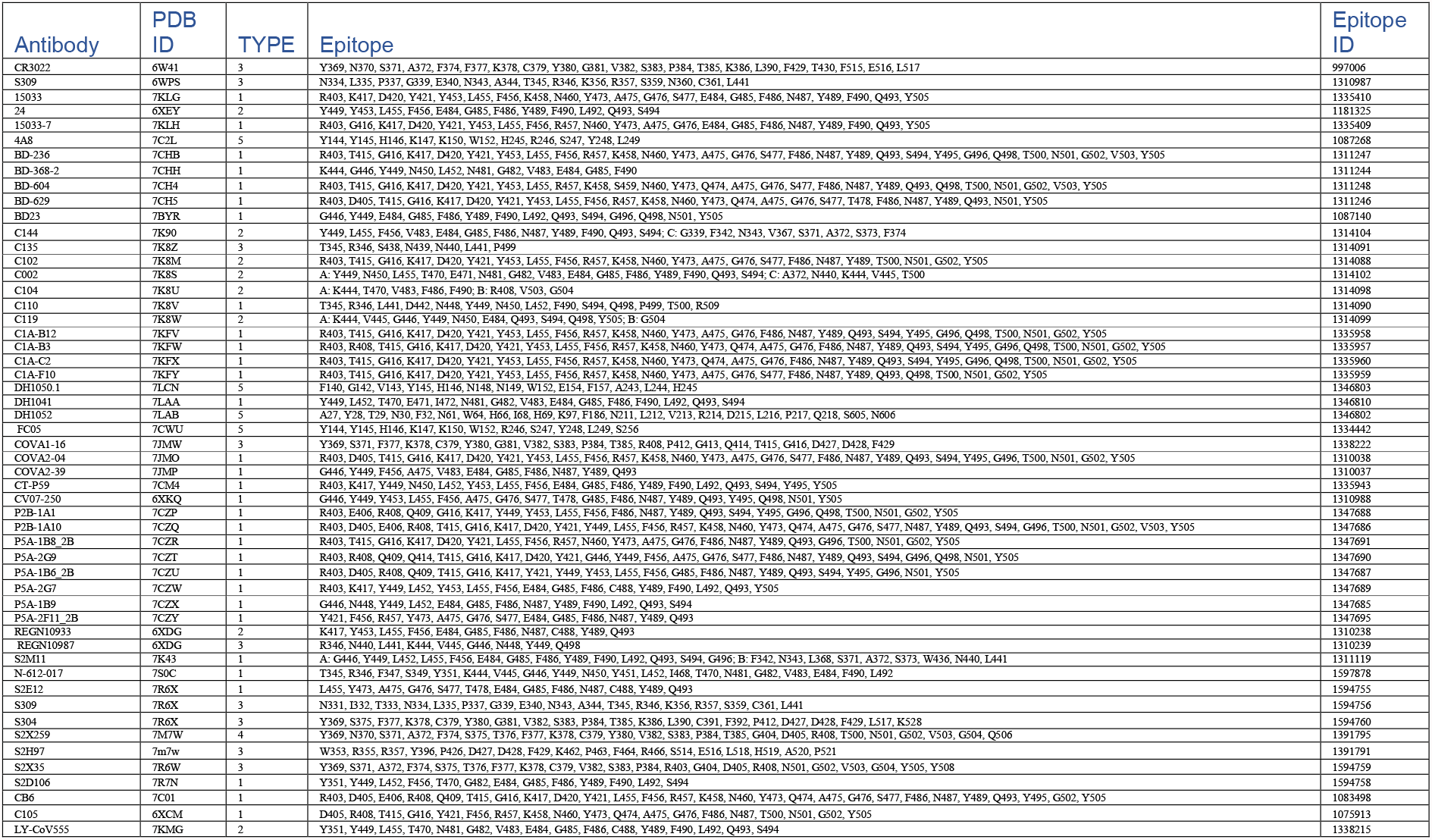
Structure and epitope details of SARS-CoV-2 spike protein targeted antibodies. Type 1-4 antibodies are RBD targeted. Type 5 are N-terminal domain antibodies. The epitope data is from IEDB database.

**Figure 2:**
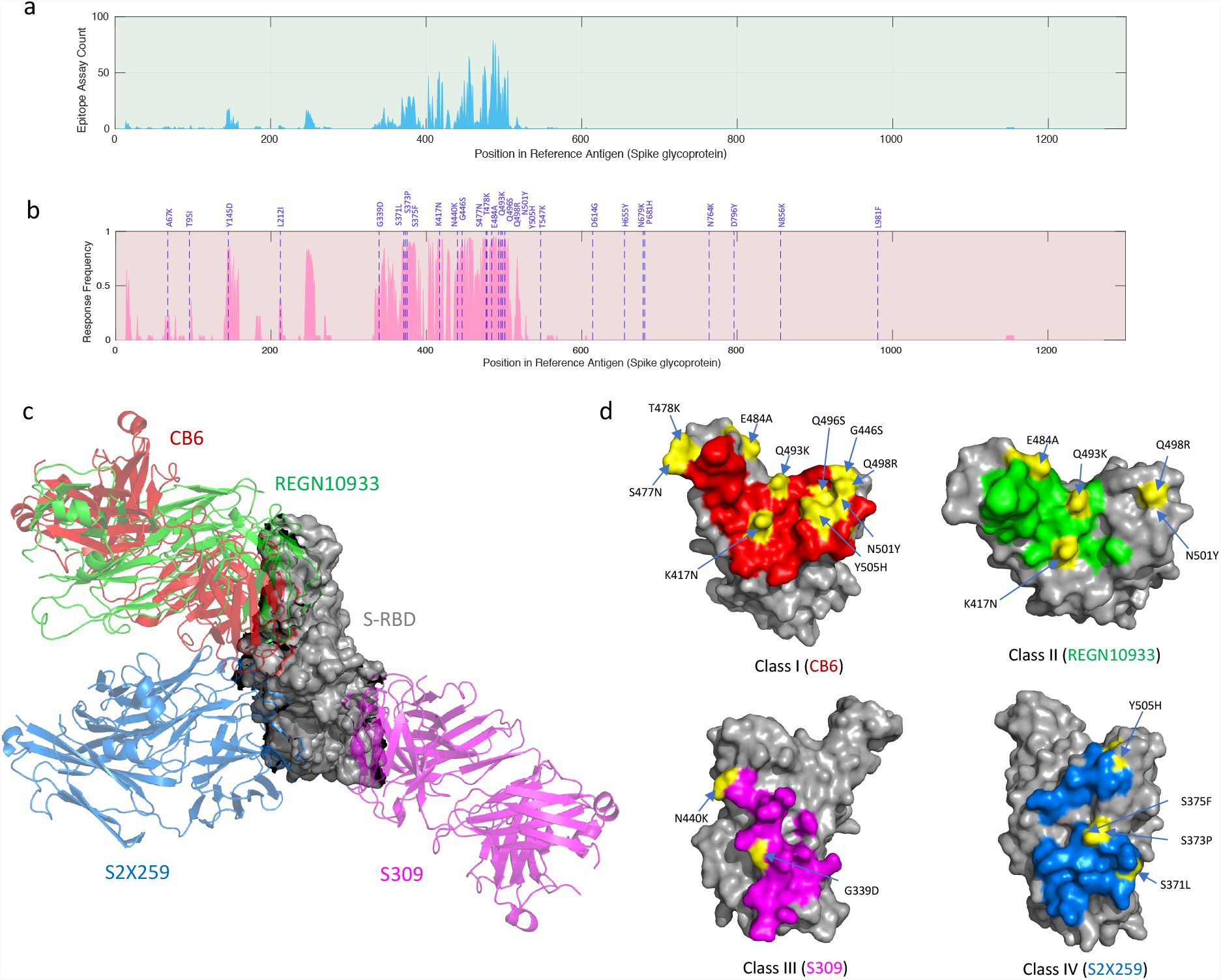
Spike protein antibodies and epitopes. (a) Graphical illustration showing number of positive epitope assays against each position in the spike protein. (b) Graphical illustration showing immune response frequency of each residue position in the spike protein. Blue lines indicate mutations in the omicron (B.1.1.529) variant. (c) RBD interaction with different antibodies. (d) Footprints of RBD targeted antibodies with the mutations in B.1.1.529 highlighted in yellow.

We selected the approved therapeutic antibodies CB6, REGN10933, and S309, along with a broadly neutralizing antibody S2X259, to understand the difference in their interaction with the WT and omicron RBD. The footprints of all four antibodies are illustrated in figure 2d with highlighted mutations found in the omicron RBD. The outcomes of the study for each antibody are discussed in the following sections.

### 3.1 Class 1: CB6 (etesevimab)

The antibody CB6, also known as etesevimab, is an approved therapeutic antibody against SARS-CoV-2. The structure of CB6 in complex with RBD (PDB ID: 7C01) and its omicron mutation-induced complex was simulated for 100 ns. MD simulation revealed the overall stability of the complexes. The root mean square deviation (RMSD) graph of the WT complex was stable; however, the backbone of the omicron mutant complex experienced fluctuations throughout the trajectory (Figure 3g). The radius of gyration (Rg) for the CB6-RBD/omicron complex deviated between 3.2 to 3.3 nm, whereas the average Rg for the CB6-RBD/WT was 3.2 nm reflecting a higher degree of compactness (Figure 3h). The minimum energy conformation of both the complexes was extracted from the trajectory for comparative interaction analysis. The total interface area between the RBD (WT) and CB6 is 1018.8 Å, which is reduced to 841.8 Å when CB6 interacts with the omicron RBD. Both the polar and non-polar interface areas are reduced for the CB6 and omicron RBD. There is a steep decline observed in the solvent accessible surface area (SASA) of the CB6-RBD/WT complex towards the end of the trajectory. An increased SASA for the omicron mutant complex may reflect conformational changes in the complex leading to a higher number of solvent exposed residues. The number of interacting residues at the interface for the WT RBD were 34; however, they were reduced to 26 in the case of omicron RBD. As a result, the total number of interface interactions also declined. The omicron mutated residues at the interface S496, R498, and Y501 fail to form hydrogen bond with CB6. Therefore, the number of hydrogen bonds were reduced from 18 to 11, resulting in an overall decrease in hydrophilic interactions at the interface.

**Figure 3:**
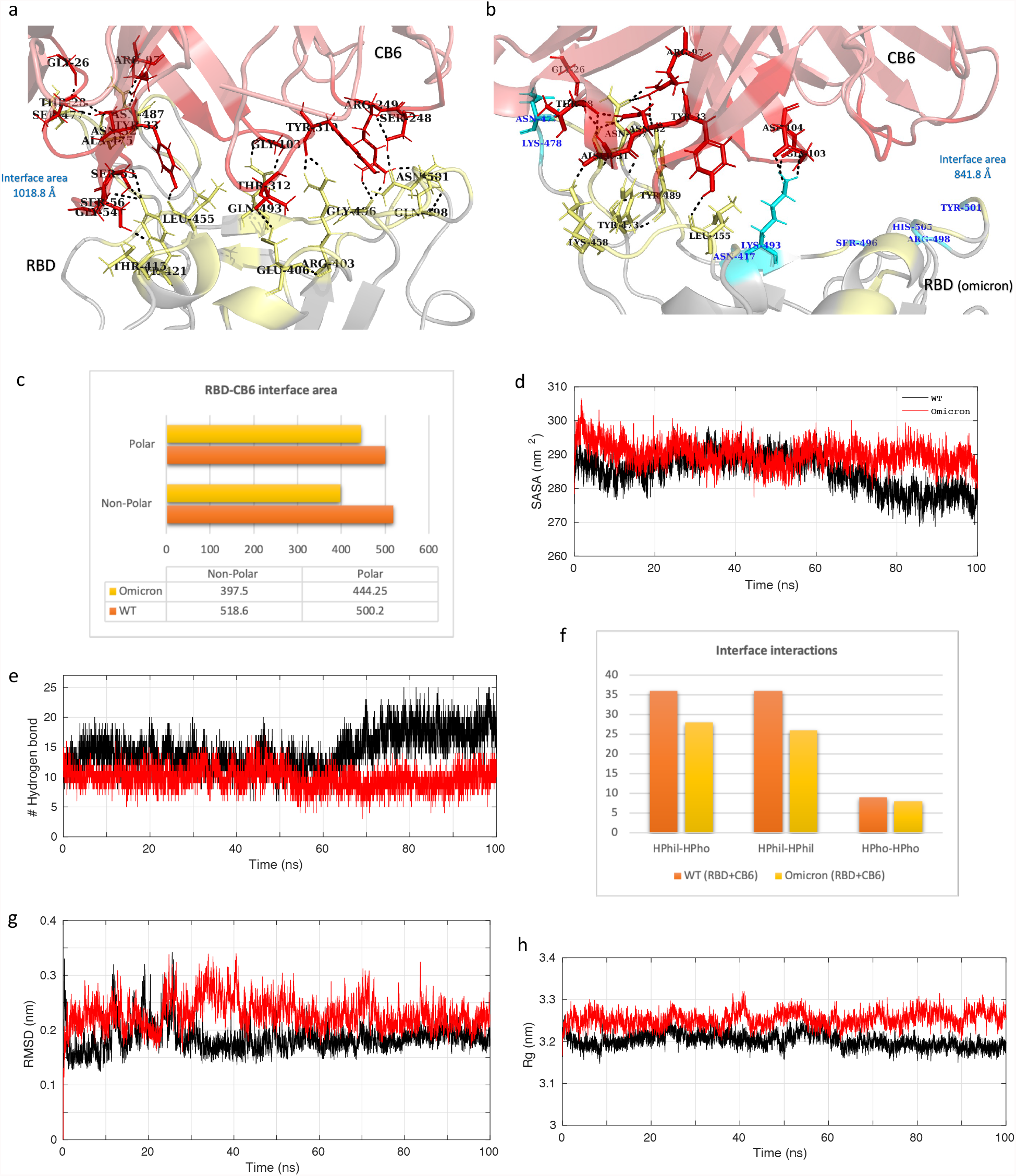
Molecular dynamics simulation results and interface statistics of RBD-CB6. (a) Wild type RBD and CB6 interface interactions. Residues involved in interactions are highlighted. Interface residues forming hydrogen bonds are labeled and shown as sticks. Hydrogen bonds are sown as black dashed lines. (b) Omicron induced mutant RBD and CB6 interface interactions. Mutant interface residues are shown in cyan and labeled in blue color. (c) Polar and nonpolar interface area of the complexes. The values are in Armstrong (Å). (d) Solvent accessible surface area (SASA) for the CB6-RBD/WT and CB6-RBD/omicron complex. (e) Plot for number of hydrogen bonds for the complexes. (f) Interface interaction count (HPhil; Hydrophilic, HPho; Hydrophobic). (g) Root mean square deviation (RMSD) plot for the complexes. (h) Radius of gyration (Rg) plot.

Further, we identified the total binding free energy and the contribution of Vander Waals, electrostatic, and solvation energies of both the complexes (Table 2). The electrostatic energy of the WT complex was −160.97 kcal/mol, which was reduced to +26.59 kcal/mol for the mutant complex. Although, there was only a slight decrease in the Vander Waals energy contribution upon mutation. However, the total binding free energy of the complexes revealed a stronger affinity between CB6 and the WT RBD than the omicron mutation-induced RBD. The residue mutations, N440K, G446S, S477N, T478K, Q493K, Q496S, Q498R, and N501Y at the CB6-RBD interface contributed negatively to the total binding free energy leading to an overall decrease in the binding affinity (Figure 4). The mutations may also affect the flexibility or rigidity of the residues at the binding interface governing the conformational dynamics and structural rearrangements of the RBD-antibody complex. The free energy landscapes (FEL) shown in figure 5 depicts the transition between various conformational ensembles of the RBD-antibody complexes. We evaluated and compared the local minima of the CB6 in complex with WT and omicron RBD. The FEL of the CB6-RBD/WT complex shows two distinct populations of conformations at the same energy basins. However, the ensembles of CB6-RBD/omicron complex have two different clusters at distinct energy basins separated by a transition barrier of 4.0 kcal/mol (Figure 5a). Hence, the omicron mutations in the RBD substantially affect the stability and the binding affinity of the CB6 antibody to its target epitope at the RBD.

**Table 2:**
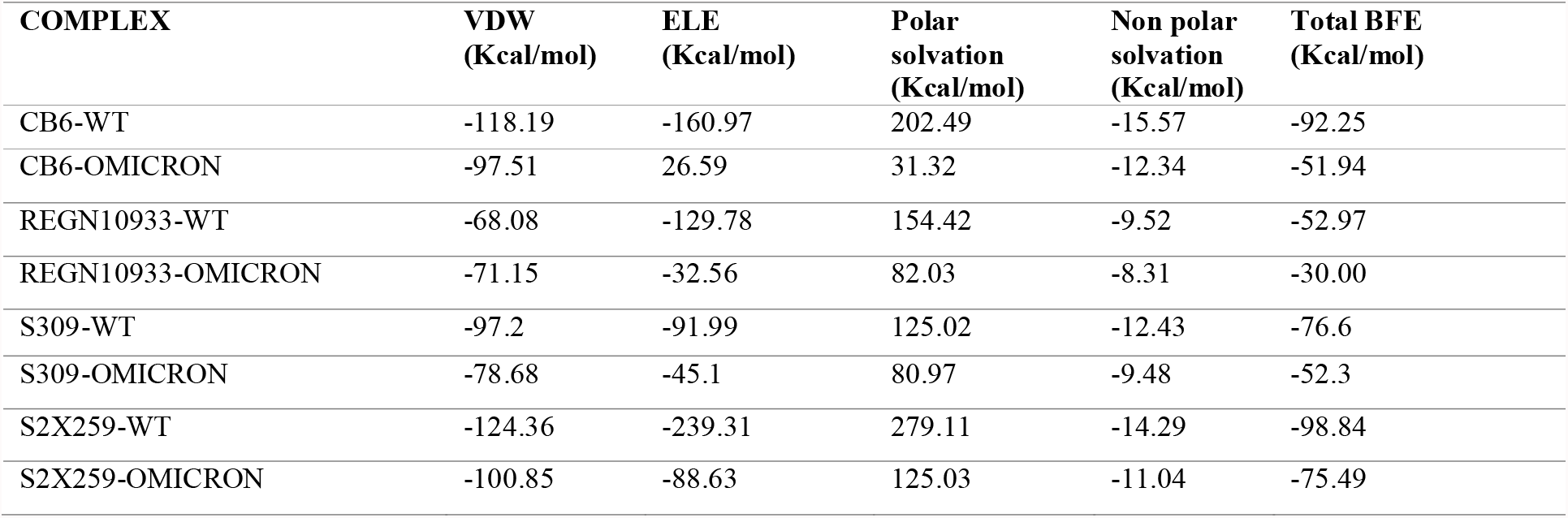
Binding free energy of the receptor binding domain and antibody complex. VDW; Vander Waals energy, ELE; electrostatic energy, BFE; Total binding free energy.

**Figure 4:**
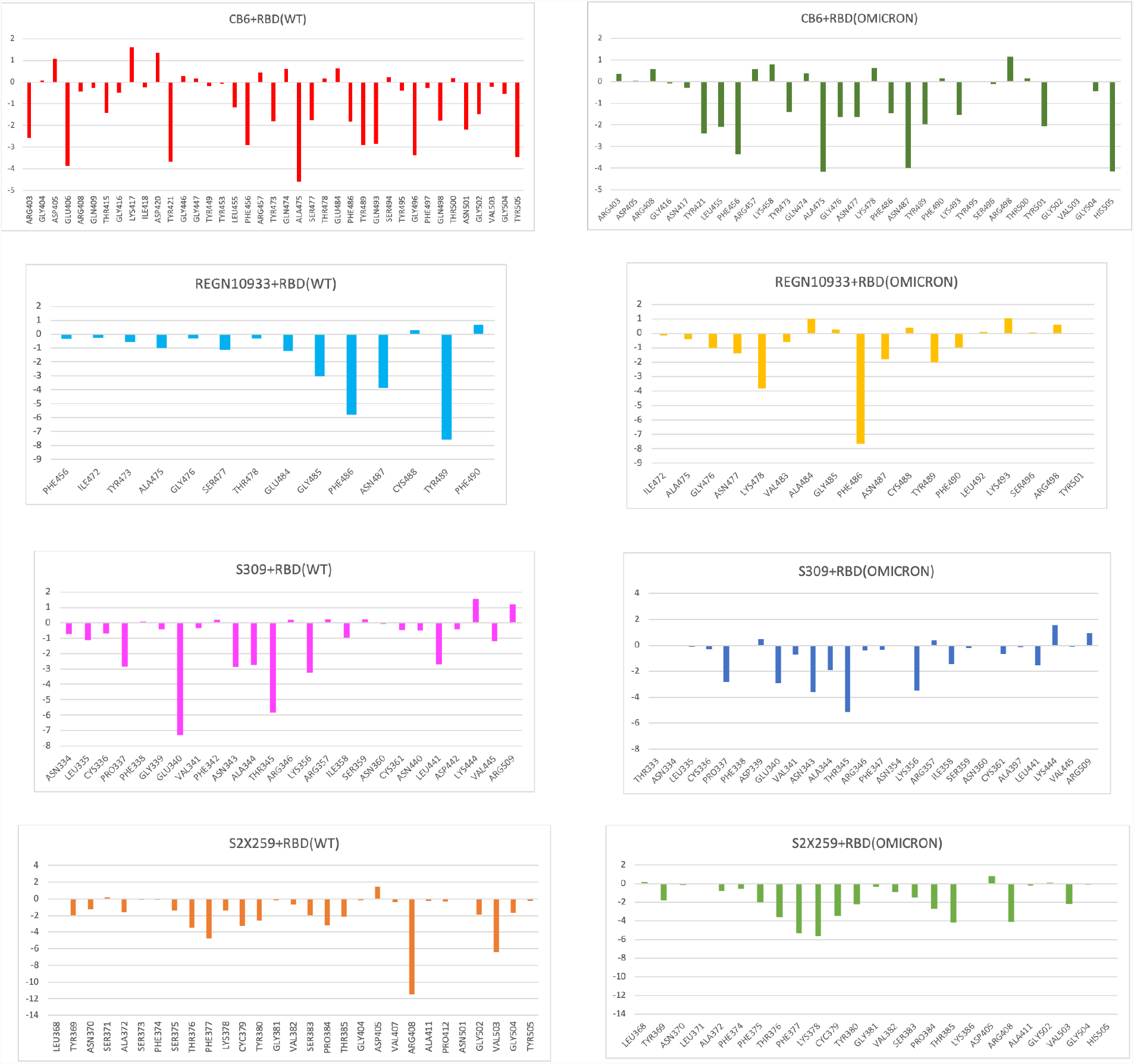
Binding free energy contribution of RBD residues involved in interaction with antibodies.

**Figure 5:**
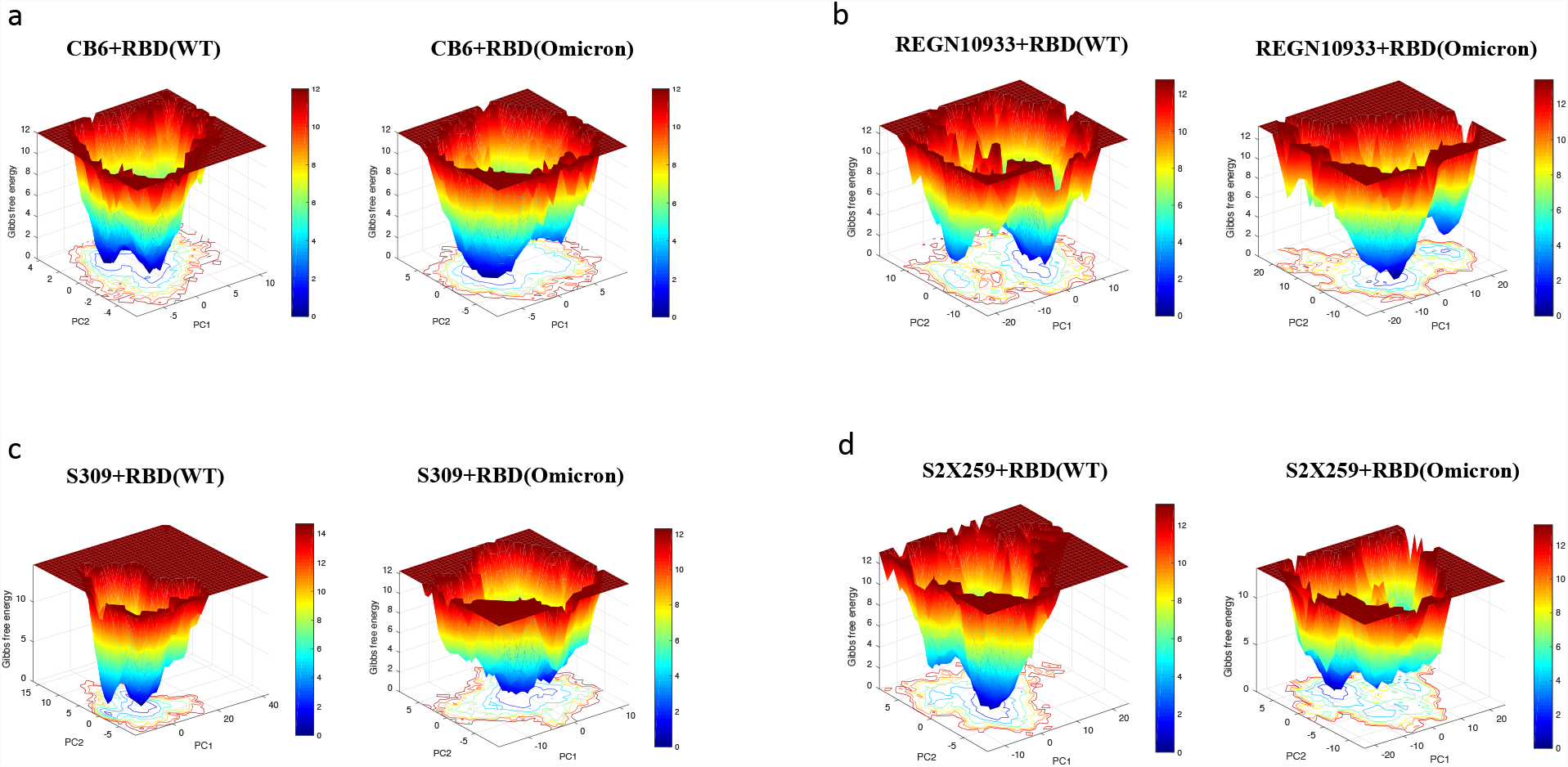
Free energy landscape plots for RBD-antibody complexes. (a) CB6 (b) REGN10933 (c) S309 (d) S2X259.

### 3.2 Class 2: REGN10933 (casirivimab)

We selected casirivimab as a class 2 antibody to evaluate the change in its binding with omicron mutation-induced RBD. REGN10933 antibody occupies the ACE2 binding region with a slightly different orientation than the CB6 antibody, as shown in figure 2c. Hence, the interface area and RBD residues involved in interaction largely differ. The antibody REGN10933 when binds to the WT RBD has an interface area of 663.25 Å, which is only reduced to 644.0 Å in the case of omicron RBD (Figure 6). Hence, no significant change was observed in the polar and non-polar interface area of the WT and mutant complexes. The SASA of these complexes remained relatively similar throughout the 100 ns MD trajectory (Figure 6e). However, the interface interaction statistics are altered upon the omicron mutations in RBD. The number of hydrophilic interactions at the interface were reduced, whereas a slight increase in the hydrophobic interactions was observed. The average number of hydrogen bonds between the antibody and RBD also reduced from 10 to 5 upon the mutations. No hydrogen bonds were formed at the interface region with a continuous stretch of mutant residues, K493, S496, R498, and Y501. Although, the overall stability of both the complexes revealed fluctuations in the backbone RMSD of both the complexes throughout the simulation trajectory. A higher degree of compactness was observed for the WT complex with Rg between 3.4 to 3.6 nm, which was observed to be greater than 3.6 nm for the omicron mutant complex.

**Figure 6:**
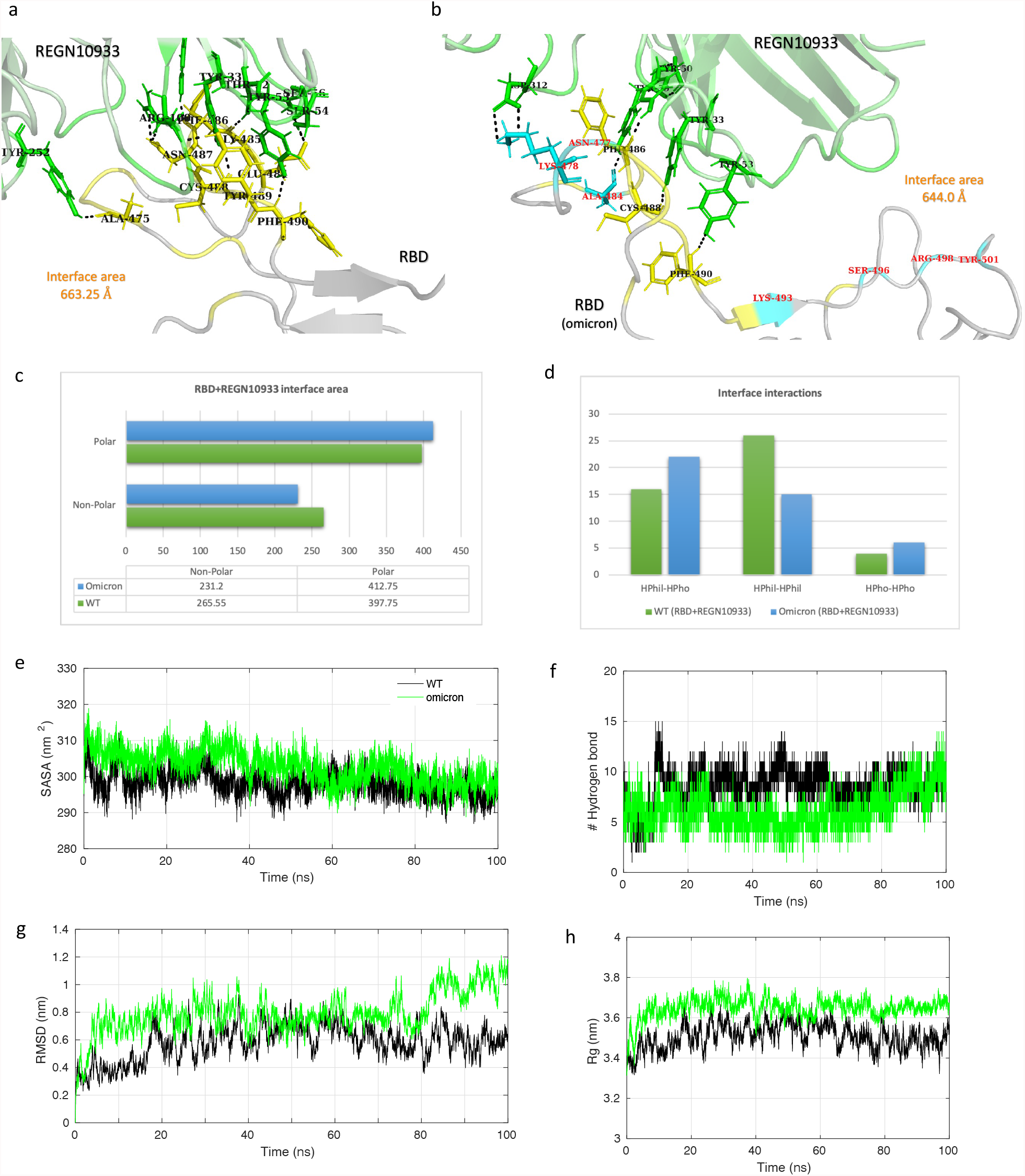
Molecular dynamics simulation results and interface statistics of RBD-REGN10933. (a) Wild type RBD and REGN10933 interface interactions. Residues involved in interactions are highlighted. Interface residues forming hydrogen bonds are labeled and shown as sticks. Hydrogen bonds are sown as black dashed lines. (b) Omicron induced mutant RBD and REGN10933 interface interactions. Mutant interface residues are shown in cyan and labeled in red color. (c) Polar and nonpolar interface area of the complexes. The values are in Armstrong (Å). (d) Interface interaction count (HPhil; Hydrophilic, HPho; Hydrophobic). (e) Solvent accessible surface area (SASA) for the REGN10933-RBD/WT and REGN10933-RBD/omicron complex. (f) Plot for number of hydrogen bonds for the complexes. (g) Root mean square deviation (RMSD) plot for the complexes. (h) Radius of gyration (Rg) plot.

The calculated binding free energy of the REGN10933-RBD/WT complex was −52.97 kcal/mol and −30.00 kcal/mol for the REGN10933-RBD/omicron complex (Table 2). This difference in the binding energy was observed due to a notable decrease in the electrostatic energy at the interface. The Vander Waals energy term for the WT and omicron mutant complex was −68.08 kcal/mol and −71.15 kcal/mol, respectively. Hence, the hydrophobic contacts at the interface were higher than the polar contacts for the mutant complex. The residue mutations S477N and T478K lead to more negative binding free energy, hence contributing positively to the total binding free energy of the complex (Figure 4). Moreover, the mutations E484A, Q493K, Q496S, and Q498R have positive values for the binding free energy, ultimately increasing the total binding free energy and hence decreasing the binding affinity of the REGN10933-RBD/omicron complex. The impact of these mutations on the lowest energy structural ensembles of each complex was observed through the FEL (Figure 5b). The FEL plot for both the complexes shows majorly two distinct conformational ensembles with a transition barrier of 2.0 kcal/mol. These observations collectively suggest that there is no significant difference in the conformational stability of REGN10933 with RBD upon omicron mutations. Although, few mutations impact the binding affinity of the RBD with the REGN10933 antibody.

### 3.3 Class 3: S309 (sotrovimab)

The NAb S309, aka sotrovimab is a therapeutic antibody approved for emergency use against the SARS-CoV-2 infection. Pre-clinical studies have suggested that sotrovimab retains activity against previously identified VOCs [36]. The antibody S309 targets an epitope in the conserved core domain of the RBD. We evaluated the binding stability and interaction of this class 3 antibody with the omicron mutation-induced RBD. The lowest energy conformation of the S309 bound to both the RBDs was extracted from the MD simulation trajectory. The total interface area between S309 and the WT RBD was 856.6 Å which reduced to 688.7 Å for the omicron RBD (Figure 7a and 7c). Along with the polar and non-polar interface area, the overall number of interactions at the interface also decreased. The average number of hydrogen bonds at the interface for the WT and omicron complex were 12 and 8, respectively (Figure 7f). The SASA for both the complexes was relatively equal up to 80 ns trajectory, and a slight decrease in the last 20 ns was observed for the WT complex. A similar trend in the trajectory was observed for the Rg of these complexes. Hence, a slightly more compact binding may occur in the case of the S309 interacting with the WT RBD. The backbone RMSD of the whole complex revealed stable conformation with lower fluctuations observed for the S309-RBD/omicron.

**Figure 7:**
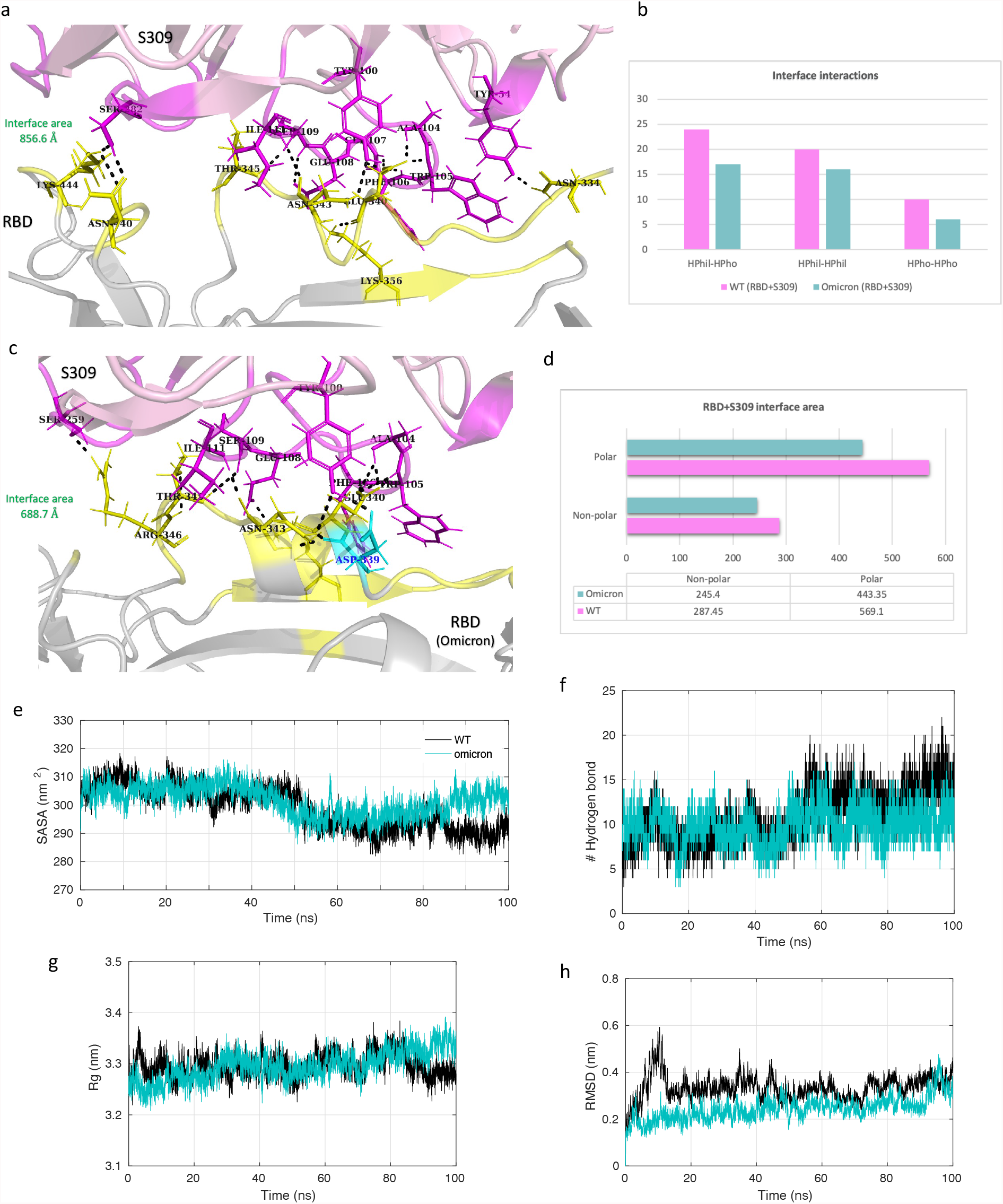
Molecular dynamics simulation results and interface statistics of RBD-S309. (a) Wild type RBD and S309 interface interactions. Residues involved in interactions are highlighted. Interface residues forming hydrogen bonds are labeled and shown as sticks. Hydrogen bonds are sown as black dashed lines. (b) Interface interaction count (HPhil; Hydrophilic, HPho; Hydrophobic). (c) Omicron induced mutant RBD and S309 interface interactions. Mutant interface residues are shown in cyan and labeled in blue color. (d) Polar and nonpolar interface area of the complexes. The values are in Armstrong (Å). (e) Solvent accessible surface area (SASA) for the S309-RBD/WT and S309-RBD/omicron complex. (f) Plot for number of hydrogen bonds for the complexes. (g) Radius of gyration (Rg) plot for the complexes. (h) Root mean square deviation (RMSD) plot.

Further, binding free energy prediction revealed the difference in the affinity of the S309 to the WT RBD and omicron mutation-induced RBD. The total binding free energy of the S309-RBD/WT and S309-RBD/omicron complex was −76.6 kcal/mol and −52.3 kcal/mol, respectively (Table 2). This free energy change was predominantly due to a decrease in the contribution of electrostatic energy in the S309-RBD/omicron complex. The residue mutation G339D leads to a positive binding free energy, contributing negatively to the total binding free energy of the complex (Figure 4). The energy contribution of the adjacent residue E340 was also reduced from −7.27 kcal/mol to −2.93 kcal/mol in the mutant complex. The binding energy and interactions were also affected by the mutation N440K. The residue N at position 440 could interact with S309 at the interface contributing its energy to the total free energy of the complex, whereas K at 440 fails to interact with S309. These changes could also be responsible for the difference in the local minima achieved by the conformation ensembles of these complexes. The FEL plot for the S309-RBD/WT complex shows that the populations of conformations are clustered in two elongated ensembles occurring at the same energy basins (Figure 5c). On the other hand, multiple populations of S309-RBD/omicron are distributed along large conformational space with a single wide cluster occurring at the local minima. Hence, these observations altogether suggest that the omicron mutations may have a limited effect on the binding of class 3 antibody S309 to the RBD.

### 3.4 Class 4: S2X259

S2X259 is a human mAb which neutralizes SARS-CoV-2, including all the previously identified VOCs, B.1.1.7, B.1.351, P.1, and B.1.427 [24]. It targets an epitope in the RBD overlapping some parts of the RBM as well as the core domain. We identified the effect of omicron mutations on this broadly neutralizing antibody. The total interface area between the RBD and the antibody was 829.85 Å, which only reduced to 812.0 Å upon mutations (Figure 8a and 8b). The SASA plot for both the complexes shows stable values throughout the 100 ns simulation, with a slightly higher area for the S2X259-RBD/omicron complex (Figure 8c). The antibody S2X259 fails to make hydrogen bond interactions near the RBM region in the omicron mutation-induced RBD (Figure 8b). Therefore, the average hydrogen bonds for the complex S2X259-RBD/WT and S2X259-RBD/omicron were 8 and 5, respectively. The residue mutation S375F resulted in the loss of hydrogen bond formation. An overall decrease in the number of hydrophilic interactions at the interface were also observed. Both the RMSD and Rg plots for the S2X259-RBD/omicron complex experienced higher fluctuation throughout the MD simulation trajectory. Hence, the mutations may affect the overall stability of the RBD-antibody complex.

**Figure 8:**
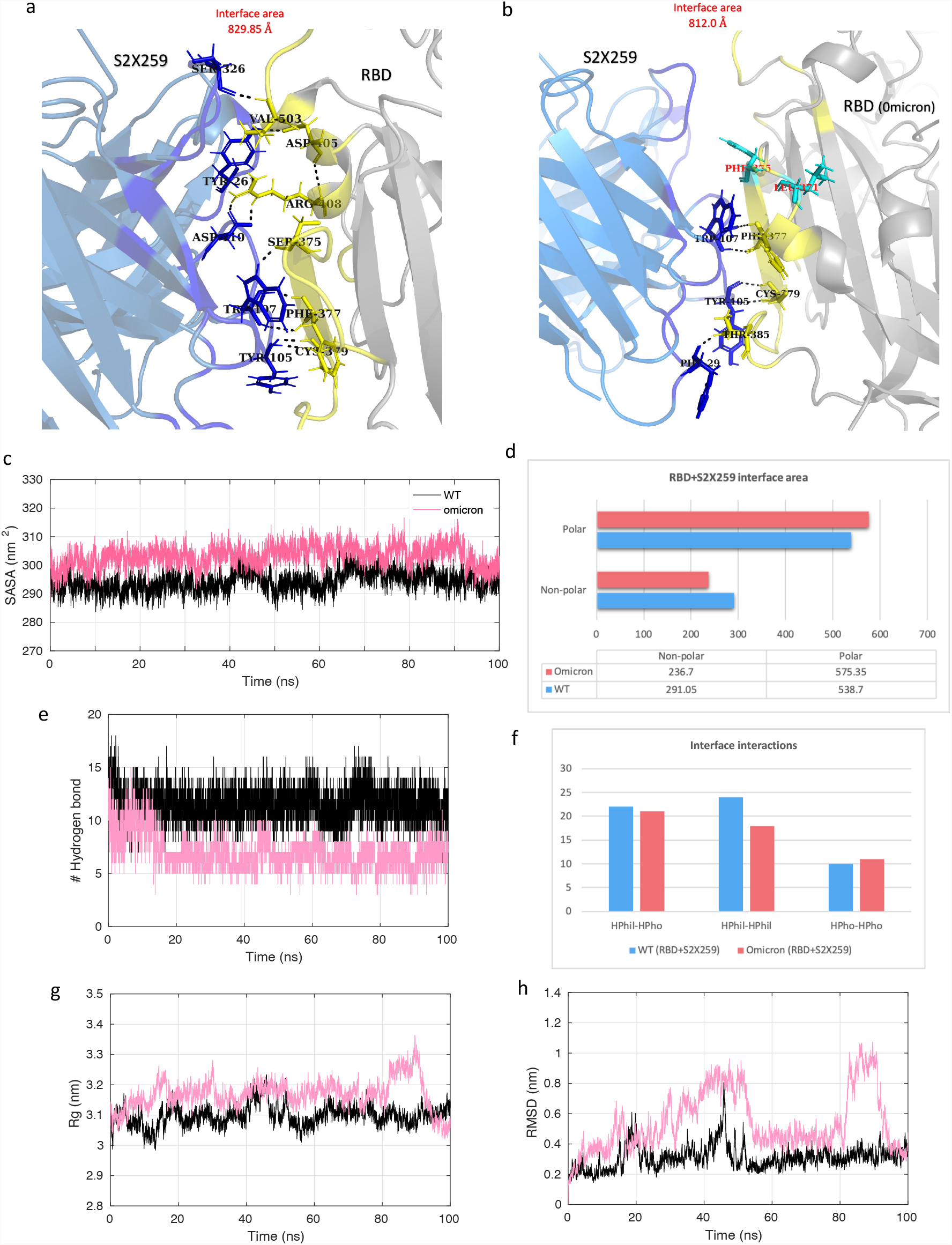
Molecular dynamics simulation results and interface statistics of RBD-S2X259. (a) Wild type RBD and S2X259 interface interactions. Residues involved in interactions are highlighted. Interface residues forming hydrogen bonds are labeled and shown as sticks. Hydrogen bonds are sown as black dashed lines. (b) Omicron induced mutant RBD and S2X259 interface interactions. Mutant interface residues are shown in cyan and labeled in red color. (c) Solvent accessible surface area (SASA) for the S2X259-RBD/WT and S2X259-RBD/omicron complex. (d) Polar and nonpolar interface area of the complexes. The values are in Armstrong (Å). (e) Plot for number of hydrogen bonds for the complexes. (f) Interface interaction count (HPhil; Hydrophilic, HPho; Hydrophobic). (g) Radius of gyration (Rg) plot for the complexes. (h) Root mean square deviation (RMSD) plot.

The difference in the binding free energy of the S2X259-RBD complexes reflects a slight decrease in the total binding affinity of the S2X259 towards the omicron RBD. Moreover, as observed for other antibodies described in the previous section, the electrostatic energy for the S2X259-RBD/omicron complex is also highly reduced (Table 2). Hence, the total binding free energy for the WT and omicron mutant complex was −98.84 kcal/mol and −75.49 kcal/mol, respectively. The residue mutations at the binding interface affect the mutant as well as adjacent residues binding energy contribution. The substitution, S371L, decreased the total binding free energy contribution of the adjacent residues N370 and A372. The residue mutations S373P and N501Y resulted in the loss of interaction with S2X259 at the interface. The substitution Y505H also contributed negatively to the total binding free energy of the S2X259 and the RBD complex (Figure 4). Further, the lowest energy conformational ensembles of the antibody and RBD complexes were evaluated through the FEL plots (Figure 5d). The FEL plot for the S2X259-RBD/WT complex shows two distinct populations of conformations confined to different energy basins separated with a transition barrier of > 4.0 kcal/mol. However, for the S2X259-RBD/omicron complex, we observed multiple ensembles at wide conformational space with energy difference >5.0 kcal/mol. Finally, the results suggest that the omicron mutations may particularly affect the binding stability of the S2X259 antibody and the RBD complex.

## 4. Conclusion

The SARS-CoV-2 omicron variant grabbed the attention as soon as it was discovered because of the large unusual set of mutations in its spike protein. The spike protein is the key antigenic determinant of the virus and an essential target for vaccine and antibody design [37]. Since the majority of therapeutic antibodies target the RBD in the spike protein, the mutations occurring in the epitope region could be of particular importance. This study reports an understanding of the structural impact of the RBD mutations on antibody binding. We evaluated four different class of RBD targeting antibodies. One therapeutic or monoclonal antibody from each class was examined to analyze the effect of omicron mutations. Class 1 antibody, CB6 aka etesevimab, acquires a maximum number of mutations in its epitope; hence, its binding affinity and stability with the RBD were substantially affected. In contrast, the antibody sotrovimab was least affected among all four antibodies by the omicron mutations. The antibody S309 or sotrovimab binds to a conserved RBD epitope with less number of mutations occurring at its interface region. However, the binding affinity of all the four antibodies was reduced for the omicron mutation-induced RBD. The mutations in the RBD largely affected and reduced the polar contacts at the interface. Hence, there was a decrease in the electrostatic energy between the antibodies and the RBD. Therefore, the omicron variant mutations occurring in the RBD may significantly reduce the efficacy of CB6, REGN10933, S309, and S2X259 antibodies. Moreover, the mutations may respond differently for different class of antibodies depending upon their target epitope and hence, their study can lead to a valuable understanding of immune escape of the virus.

## Acknowledgment

We acknowledge the financial assistance provided by the University Grant Commission in the form of Junior Research Fellowship (JRF) to Jyoti Verma.

## Reference

1. Classification of Omicron (B.1.1.529): SARS-CoV-2 Variant of Concern. https://www.who.int/news/item/26-11-2021-classification-of-omicron-(b.1.1.529)-sars-cov-2-variant-of-concern. Accessed 19 Jan 2022

2. Wolter N, Jassat W, Walaza S, et al (2021) Early assessment of the clinical severity of the SARS-CoV-2 Omicron variant in South Africa. medRxiv 2021.12.21.21268116. https://doi.org/10.1101/2021.12.21.21268116

3. GISAID - hCov19 Variants. https://www.gisaid.org/hcov19-variants/. Accessed January 25 2022

4. CoVariants. https://covariants.org/variants/21K.Omicron. Accessed January 19 2022

5. Gong SY, Chatterjee D, Richard J, et al (2021) Contribution of single mutations to selected SARS-CoV-2 emerging variants spike antigenicity. Virology 563:134–145. https://doi.org/10.1016/J.VIROL.2021.09.001

6. Zahradník J, Marciano S, Shemesh M, et al (2021) SARS-CoV-2 variant prediction and antiviral drug design are enabled by RBD in vitro evolution. Nat Microbiol 2021 69 6:1188–1198. https://doi.org/10.1038/s41564-021-00954-4

7. Verma J, Subbarao N (2021) Insilico study on the effect of SARS-CoV-2 RBD hotspot mutants’ interaction with ACE2 to understand the binding affinity and stability. Virology 561:107–116. https://doi.org/10.1016/J.VIROL.2021.06.009

8. Harvey WT, Carabelli AM, Jackson B, et al (2021) SARS-CoV-2 variants, spike mutations and immune escape. Nat Rev Microbiol 2021 197 19:409–424. https://doi.org/10.1038/s41579-021-00573-0

9. Wang P, Nair MS, Liu L, et al (2021) Antibody resistance of SARS-CoV-2 variants B.1.351 and B.1.1.7. Nat 2021 5937857 593:130–135. https://doi.org/10.1038/s41586-021-03398-2

10. Wilhelm A, Widera M, Grikscheit K, et al (2021) Reduced Neutralization of SARS-CoV-2 Omicron Variant by Vaccine Sera and monoclonal antibodies. medRxiv 2021.12.07.21267432. https://doi.org/10.1101/2021.12.07.21267432

11. Liu L, Iketani S, Guo Y, et al (2021) Striking Antibody Evasion Manifested by the Omicron Variant of SARS-CoV-2. https://doi.org/10.1038/s41586-021-04388-0

12. Cao Y, Wang J, Jian F, et al Omicron escapes the majority of existing SARS-CoV-2 neutralizing antibodies. https://doi.org/10.1038/s41586-021-04385-3

13. Chen J, Wang R, Gilby NB, Wei G-W (2021) Omicron (B.1.1.529): Infectivity, vaccine breakthrough, and antibody resistance. ArXiv

14. Barnes CO, Jette CA, Abernathy ME, et al (2020) SARS-CoV-2 neutralizing antibody structures inform therapeutic strategies. Nat 2020 5887839 588:682–687. https://doi.org/10.1038/s41586-020-2852-1

15. Shi R, Shan C, Duan X, et al (2020) A human neutralizing antibody targets the receptor-binding site of SARS-CoV-2. Nat 2020 5847819 584:120–124. https://doi.org/10.1038/s41586-020-2381-y

16. Ju B, Zhang Q, Ge J, et al (2020) Human neutralizing antibodies elicited by SARS-CoV-2 infection. Nat 2020 5847819 584:115–119. https://doi.org/10.1038/s41586-020-2380-z

17. Liu L, Wang P, Nair MS, et al (2020) Potent neutralizing antibodies against multiple epitopes on SARS-CoV-2 spike. Nat 2020 5847821 584:450–456. https://doi.org/10.1038/s41586-020-2571-7

18. Hansen J, Baum A, Pascal KE, et al (2020) Studies in humanized mice and convalescent humans yield a SARS-CoV-2 antibody cocktail. Science (80-) 369:1010–1014. https://doi.org/10.1126/SCIENCE.ABD0827/SUPPL_FILE/ABD0827_MDAR-REPRODUCIBILITYCHECKLIST.PDF

19. Zost SJ, Gilchuk P, Case JB, et al (2020) Potently neutralizing and protective human antibodies against SARS-CoV-2. Nat 2020 5847821 584:443–449. https://doi.org/10.1038/s41586-020-2548-6

20. Jones BE, Brown-Augsburger PL, Corbett KS, et al (2021) The neutralizing antibody, LY-CoV555, protects against SARS-CoV-2 infection in nonhuman primates. Sci Transl Med 13:p. https://doi.org/10.1126/SCITRANSLMED.ABF1906/SUPPL_FILE/ABF1906_SM.PD F

21. Pinto D, Park YJ, Beltramello M, et al (2020) Cross-neutralization of SARS-CoV-2 by a human monoclonal SARS-CoV antibody. Nat 2020 5837815 583:290–295. https://doi.org/10.1038/s41586-020-2349-y

22. Li D, Edwards RJ, Manne K, et al (2021) In vitro and in vivo functions of SARS-CoV-2 infection-enhancing and neutralizing antibodies. Cell 184:4203–4219.e32. https://doi.org/10.1016/J.CELL.2021.06.021

23. Garrett Rappazzo C, Tse L V., Kaku CI, et al (2021) Broad and potent activity against SARS-like viruses by an engineered human monoclonal antibody. Science (80-) 371:823–829. https://doi.org/10.1126/SCIENCE.ABF4830/SUPPL_FILE/ABF4830_TABLE_S2_GISAID_ACKNOWLEDGMENTS.PDF

24. Tortorici MA, Czudnochowski N, Starr TN, et al (2021) Broad sarbecovirus neutralization by a human monoclonal antibody. Nat 2021 5977874 597:103–108. https://doi.org/10.1038/s41586-021-03817-4

25. surface glycoprotein [Severe acute respiratory syndrome coronavirus 2] -Protein - NCBI. https://www.ncbi.nlm.nih.gov/protein/YP_009724390.1/. Accessed January 19 2022

26. surface glycoprotein [Severe acute respiratory syndrome coronavirus 2] -Protein -NCBI. https://www.ncbi.nlm.nih.gov/protein/UFO69279.1. Accessed January 19 2022

27. EMBOSS Needle < Pairwise Sequence Alignment < EMBL-EBI. https://www.ebi.ac.uk/Tools/psa/emboss_needle/. Accessed January 19 2022

28. Berman HM, Westbrook J, Feng Z, et al (2000) The Protein Data Bank. Nucleic Acids Res 28:235–242. https://doi.org/10.1093/NAR/28.1.235

29. IEDB.org: Free epitope database and prediction resource. https://www.iedb.org/. Accessed January 19 2022

30. Maestro | Schrödinger

31. Berendsen HJC, Van Der Spoel D, Van Drunen R (1995) GROMACS: A message-passing parallel molecular dynamics implementation. Comput Phys Commun 91:43–56

32. CoCoMaps Tool. https://www.molnac.unisa.it/BioTools/cocomaps/. Accessed January 24 2022

33. Vangone A, Spinelli R, Scarano V, et al (2011) COCOMAPS: a web application to analyze and visualize contacts at the interface of biomolecular complexes. Bioinformatics 27:2915–2916. https://doi.org/10.1093/BIOINFORMATICS/BTR484

34. Weng G, Wang E, Wang Z, et al (2019) HawkDock: a web server to predict and analyze the protein–protein complex based on computational docking and MM/GBSA. Nucleic Acids Res 47:W322–W330. https://doi.org/10.1093/NAR/GKZ397

35. HawkDock Server. http://cadd.zju.edu.cn/hawkdock/#MMGBSA_submit. Accessed January 24 2022

36. Chen RE, Winkler ES, Case JB, et al (2021) In vivo monoclonal antibody efficacy against SARS-CoV-2 variant strains. Nat 2021 5967870 596:103–108. https://doi.org/10.1038/s41586-021-03720-y

37. Walls AC, Park Y-J, Tortorici MA, et al (2020) Structure, Function, and Antigenicity of the SARS-CoV-2 Spike Glycoprotein. Cell 181:281–292.e6. https://doi.org/10.1016/j.cell.2020.02.058

